# Directional selection at gene expression level contributes to the speciation of Asian rice cultivars

**DOI:** 10.1101/2021.10.31.466501

**Authors:** Lihong Xie, Kehan Yu, Dongjing Chen

## Abstract

Differences in expression levels play important roles in phenotypic variation across species, especially those closely related species with limited genomic differences. Therefore, studying gene evolution at expression level is important for illustrating phenotypic differentiation between species, such as the two Asian rice cultivars, *Oryza sativa* L. ssp. *indica* and *Oryza sativa* L. ssp. *japonica*. In this study, we evaluated the gene expression variation at inter-subspecies and intra-subspecies level using transcriptome data from seedlings of three *indica* and *japonica* rice and defined four groups of genes under different natural selections. We found a substantial of genes (about 79%) that are under stabilizing selection at the expression level in both subspecies, while about 16% of genes are under directional selection. Genes under directional selection have higher expression level and lower expression variation than those under stabilizing selection, which suggest a potential explanation to subspecies adaptation to different environments and interspecific phenotypic differences. Subsequent functional enrichment analysis of genes under directional selection shows that *indica* rice have experienced the adaptation to environmental stresses, and also show differences in biosynthesis and metabolism pathways. Our study provide an avenue of investigating *indica*-*japonica* differentiation through gene expression variation, which may guide to rice breeding and yield improvement.

## Introduction

Rice grown in Asia (*Oryza sativa* L.) is a very important food crop for China and the world (Ruan Bosheng, 2008), and more than half of the world’s population uses it as a staple food (Sasaki & Burr, 2000). It is mainly divided into two different subspecies, namely the subspecies *Oryza sativa* L. ssp. *indica* (*indica*) and the subspecies *Oryza sativa* L. ssp. *japonica* (*japonica*). It is speculated that *japonica* and *indica* rice were domesticated 9,000 years ago (Purugganan & Fuller, 2009), but debate about their origins still exists. The first model, the single-source model, indicated that the two main subspecies of Asian rice, *indica* and *japonica*, were domesticated from wild rice (*O. rufipogon.*) and then differentiated (Ting, Y., 1957) (Lu et al. al., 2002; Wang et al., 2008). In contrast, the second model, the multiple independent domestication model, proposes that the two main rice types are domesticated separately (Oka & Morishima, 1982). *Indica* rice evolved from wild rice. Because, *japonica* rice is the continuous evolution and artificial selection of *indica* rice in the process of people’s continuous introduction to high latitudes and high altitude areas (Lu Baorong et al., 2009). Therefore, *indica* rice is the basic type, and *japonica* rice is the variant type (Molina et al., 2011). The latter has been supported by many researchers after the observation of the strong genetic differentiation between *indica* and *japonica* and the development of several systems of rice domestication (Wang, X. et al., 1984). The third origin model claims that *indica* and *japonica* were independently domesticated or at least domesticated twice and then differentiated from wild rice (*O. rufipogon*.) (Garris et al., 2005; Second, 1982).

In most cases, *indica* rice ecotypes are mainly distributed in low-latitude or high-altitude tropical and subtropical rice growing areas, while *japonica* rice ecotypes are mainly distributed in high-latitude temperate areas (Lu et al., 2009). Due to long-term adaptation to different ecological environments, *japonica* rice and *indica* rice have differences in morphological characteristics, agronomic traits, and genes. In terms of morphological characteristics, the stems of *indica* rice are thicker, and the plant height is generally more than 1 meter. The tillering ability is stronger, the leaf color is lighter, the grains are slender, easy to fall, and the rice yield rate is low. However, *japonica* rice generally has a thin stem and a plant height of 75-95 cm. Traditional *japonica* rice varieties have a lower tillering ability than *indica* rice. The leaves are darker, the grains are short and round, and they are not easy to shatter, and the rice yield rate is higher. (*Flower Encyclopedia;* Xu Zhengjin et al., 2003). In terms of agronomic characteristics, *indica* rice with a short growth period is more resistant to humidity, heat, and strong light, but not cold tolerant. After being hulled into *indica* rice, the transparency of the rice grains is low. Because it contains about 20% amylose, *indica* rice is drier and looser when cooked. However, *japonica* rice, which has a long growth period, generally only matures once a year, is more cold-tolerant and tolerant to low light, but not tolerant to high temperatures. After being hulled into *japonica* rice, the rice grains have high transparency(https://Douding.com). Because it contains less amylose, less than 15%, it is medium-viscosity (Report on Rice Factory Seedling https://Breeding-Douding.com), and its cooked food characteristics are between glutinous rice and *indica* rice (Jiang Jian et al., 2001; Miao Xiangwei & Wang Dexin, 2009; Xu Hai et al., 2007).

We want to study how the advantages of gene expression evolution between *japonica* rice and *indica* rice are reflected in the phenotype. *Japonica* rice and *indica* rice are two subspecies with similar evolutionary distances and similar genomes. Because the differences between species caused by DNA sequences are relatively small, the differences caused by gene expression regulation are very important for the interpretation of phenotypic differences.

Previous studies on phenotypic differences in species focused on the evolution of gene lineage to study species differentiation (Onishi et al., 2007; Ting et al., 2000), but a small number of study focused on the study of transcriptomes. The transcriptome is a collection of all transcription products in a cell under a specific developmental stage or physiological condition of a species. Understanding the transcriptome is of great significance for explaining the functional elements of the genome and the phenotypic traits controlled by its line (Wang et al., 2009). Therefore, we want to study the influence of the transcriptome on the differences in gene expression and the changes in traits. This will provide great research significance for species differentiation.

In order to measure the extent and specific traits of the differences between Asian cultivated rice subspecies caused by gene expression levels, we divided genes into four categories according to the differences in gene expression between subspecies and within subspecies: (1) genes subjected to directional selection: genes with large differences in expression between species and small differences in expression within species are the genes that continue to strengthen the selection pressure for a certain trait among populations. These genes are also the main genes that cause differences in traits between species (Mitchell-Olds et al., 2007; Rieseberg et al., 2002). (2) Genes subjected to stabilizing selection: genes with small differences between species and within species are the continuous selection pressure for interspecies traits in the population. This natural selection will not lead to population differentiation (Charlesworth et al., 1982). (3) Genetic drift: genes with large differences between species and within species are a type of gene affected by random factors in the population (Melo & Marroig, 2015; Nielsen, 2005). (4) Complex scenario: genes with small inter-species differences and large intra-species differences. After the identification of natural selection, we analyze for gene enrichment of the genes subject to directional selection and understand its specific traits and functions (Subramanian et al., 2005).

This article studies the following scientific issues. First, whether the genes subject to directional selection in expression are the genes that cause the phenotypic differences between *japonica* rice and *indica* rice. Second, whether the functions of these genes are related to phenotypic differences.

The purpose and significance of this research are mainly to increase yield and breeding. It is to study that this species is closely related to the food production of all human beings, and to analyze the reasons for the differences in rice phenotypes so that people can better control the traits to achieve the ideal yield and quality of rice.

## Materials and methods

### The acquisition of rice transcriptome data and the measurement of gene expression levels

We obtained the seedling transcriptome data of three different *indica* and *japonica* rice lines (*Indica*: Khao Dawk Mali 105, Guangluai 4, and Rathuwee; *Japonica*: Taipei 309, Heukgyeong, and Nipponbare) from Wen et al. (Wen et al., 2016). Subsequently, we used hisat2 (version: 2.2.0, default parameters) to map the read length of the original sequencing data of all samples back to the *japonica* rice reference genome (ensembl, IRGSP-1.0), and used stringtie (version: 2.1.6; parameters: -e, Other defaults) to calculate the TPM (transcript per million) of the protein-coding gene (ensembl, IRGSP-1.0.51). TPM was used as the expression level of the protein-coding gene for subsequent analysis.

### Principal component analysis

After obtaining gene expression levels, we use principal components to analyze the differences between samples. First, the gene-specific TPM uses the R method vegdist to calculate the dissimilarity index between samples, then uses the R method prcomp to calculate the principal component of the sample for the index, and finally uses the R method ggbiplot to visualize the data. (Figure 1, grouping, principle component separates different samples)

**Figure 1.**
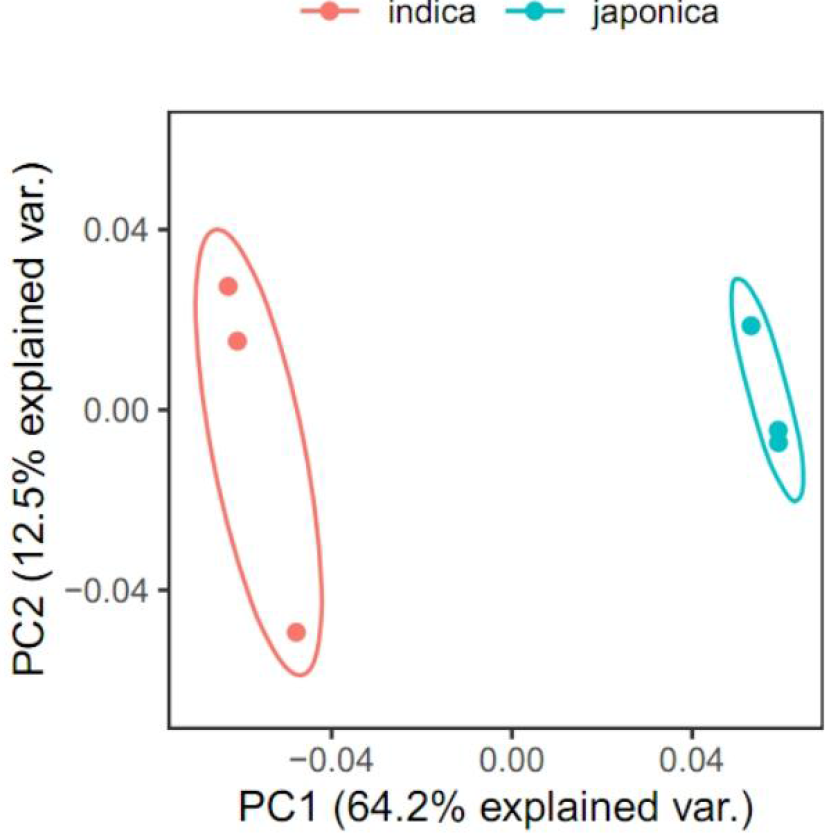
Principal component analysis of the transcriptome of *indica* and *japonica* rice samples.

### Identification of genes under four types of natural selection

We adopted the method of Yeh et al. According to whether there are significant differences in gene expression between species and within species, we divided them into four categories: directional selection: significant differences between species and insignificant differences within species; stabilizing selection: both difference between interspecies and difference between intraspecies are not significant; genetic drift: interspecies and intraspecies differences are both significant; complex scenario: interspecies differences are not significant, and intraspecies differences are significant (Yeh et al., 2014). We use a linear model to measure whether there are significant differences in gene expression between species and within species.

For interspecies:

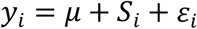

Among them, *y*_*i*_ represents the expression level of the protein coding gene of species i, μ is the basic expression level, *S*_*i*_ is the effect of species i, and *ε*_*i*_ is the residual. If the P value of the linear model is less than 0.05, it is defined as significant difference between species.

For intraspecies:

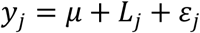

Among them, *y*_*j*_ represents the expression level of the protein coding gene of line j, μ is the basic expression level, *L*_*j*_ is the effect of line j, *ε*_*j*_ and is the residual. If the P value of the linear model is less than 0.05, it is defined as significant intraspecies difference.

### Functional enrichment analysis

In order to find genes and functions related to subspecies differentiation, we used RiceNETDB (http://bis.zju.edu.cn/ricenetdb/) to carry out Gene Ontology (Gene Ontology) on *indica* and *japonica* species-specific directional selection genes. , GO) enrichment analysis. The three functional branches of GO, the biological process, molecular function and cellular component, are all used for functional enrichment analysis.

## Results

### Processing and expression analysis of Asian cultivated rice transcriptome data

We obtained the seedling transcriptome data of three *indica* rice lines (Khao Dawk Mali 105, Guangluai 4 and Rathuwee) and three *japonica* rice lines (Taipei 309, Heukgyeong and Nipponbare) from the research of Wen et al. (Wen et al., 2016)(Table 1). After hisat2 (version 2.2.0), the initial read length was posted to the reference genome (ensembl, IRGSP-1.0). Among the samples with a total read length of 18,818,192 to 20,326,850, it was found that the only response rate of the transcriptome sequencing read length was between 63.41 and 68.40%. For the 35,775 protein-coding genes that have been annotated, the number of genes detected to be expressed (TPM greater than 0) ranged from 28,021 to 29,092 (78.33 to 81.32%); while higher expression levels were detected (TPM greater than or equal to 5), the number of genes is between 16,942-17,807 (47.36-49.77%). It shows that in our subsequent analysis, about 50% of the genes can be judged to be credible and selected because of their higher expression levels.

**Table 1.** *Indica* and *Japonica* transcriptome statistics

| Subspecies | SRR id | Accession name | Total reads | Uniquely mapped | # detected PCGs (TPM>0) | # of # detected PCGs (TPM≥5) |
| --- | --- | --- | --- | --- | --- | --- |
| *indica* | SRR2154082 | Guangluai4 | 20,326,850 | 13,336,411 (65.61%) | 28,021 (78.33%) | 16,942 (47.36%) |
|  | SRR2154083 | KDM105 | 20,216,686 | 12,820,328 (63.41%) | 28,316 (79.15%) | 17,367 (48.55%) |
|  | SRR2154084 | Rathuwee | 19,876,912 | 12,716,807 (63.98%) | 28,240 (78.94%) | 17,428 (48.72%) |
| *japonica* | SRR2154085 | Heukgyeong | 19,889,198 | 13,584,731 (68.30%) | 28,766 (80.41%) | 17,312 (48.39%) |
|  | SRR2154086 | Nipponbare | 18,818,192 | 12,871,620 (68.40%) | 29,300 (81.90%) | 17,807 (49.77%) |
|  | SRR2154087 | Taipei309 | 19,094,551 | 13,013,267 (68.15%) | 29,092 (81.32%) | 17,475 (48.85%) |

Subsequently, we conducted a principal component analysis of six lines from two cultivated rice subspecies, and found that the difference between the samples of *indica* and *japonica* is relatively large (the first principal component PC1 can explain 64.2% of the variation between the samples), while the subspecies differences in the internal samples are small (the second principal component PC2 only explains 12.5% of the variation between samples) (Figure 1). It shows that the data is sufficient to define the evolution of expression based on inter-species and intra-species differences (see Materials and Methods).

### The expression of most genes in *indica* and *japonica* rice is affected by stabilizing selection

*Indica* rice and *japonica* rice are subspecies of cultivated rice in Asia. They diverged about 0.55 million years ago (Stein et al. 2018, nature genetics) . There is only 1/6 difference in genome , and most of them are located on transposons (Campbell et al., 2020; Gao et al., 2015; Ma & Bennetzen, 2004). Therefore, in addition to the genomic differences between *indica* and *japonica* rice, differences in gene expression levels are also an important factor explaining the differences in traits between this subspecies (Rieseberg et al., 2002). In order to find out which genes cause differences between subspecies, we divided genes into four categories based on differences in gene expression between and within subspecies: genes subject to directional selection, genes subject to stabilizing selection gene, genetic drift, and complex scenario. Our results show that among the 35,775 genes in *indica* and *japonica*, most of the genes are subject to stabilizing selection (*indica*: 28,501, accounting for 79.67%; *japonica*: 28,198 , accounting for 78.82%), which is similar to the two subspecies. The genomic differences are relatively small and consistent (Figure 2A,B). For example, the expression difference of the gene Os12g0274700 between and within the two subspecies is relatively small and not significant (Figure 2C, D). However, there are still quite a few genes subject to targeted selection, 16.25% (5,910/35,775) in *indica* rice and 16.36% ( 5,854/35,775) in *japonica* rice (Figure 1A, B). For example, in *indica* rice, the gene Os08g0435900 , which is subject to targeted selection, has a large difference in expression between species, while the difference between *indica* species is not significant (Figure 2C). This gene is chlorophyll ab binding protein P4 (chlorophyll ab binding protein P4), which is located in the chloroplast, and is highly expressed in flag leaves before flowering (Wang et al., 2015) . The gene Os07g0147500 , which is subject to directional selection in *japonica* rice, has a larger expression difference between *japonica* rice and *indica* rice, but the expression difference in *japonica* rice is not significant (Figure 2D). This gene is a 10kDa polypeptide located in the photosystem II of the chloroplast , and is highly expressed in flower buds, leaves after anthesis, and grain-filled seeds (Wang et al., 2015) . In addition, genetic drift genes whose expression is affected by random factors, that is, genes that are not significantly different between species and within species, account for about 3.09~3.94% of the two subspecies; the expression of very few genes differs within species Significant but not significant differences between species, it is a complex situation, accounting for 0.72 to 0.88% (Figure 2A, B).

**Figure 2.**
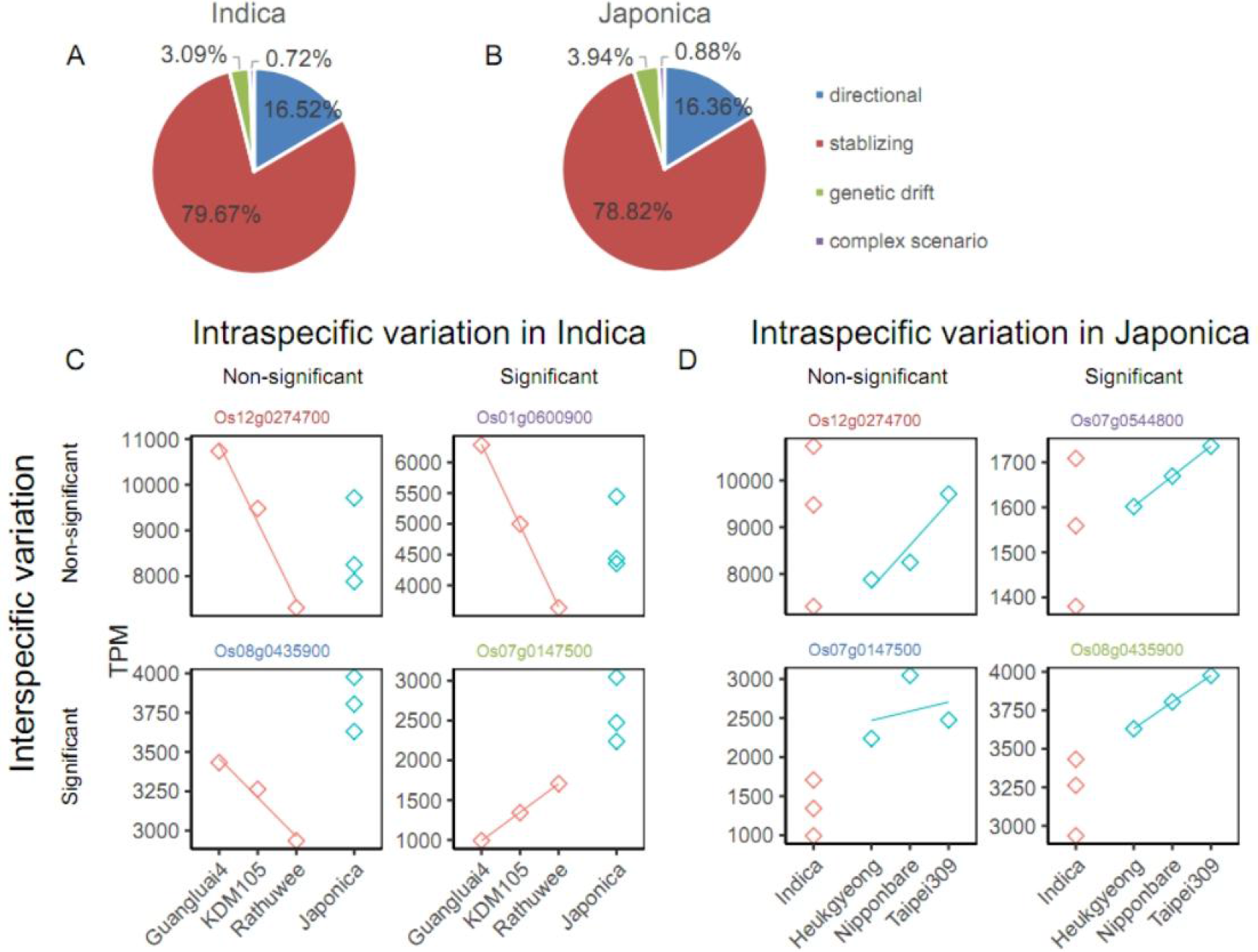
Evolutionary pattern of gene expression in *indica* and *japonica* rice. The ratio of the four types of expression selection in *indica* (A) and *japonica* (B). Graphical representation of the expression of four different selection types of genes in *indica* (C) and *japonica* (D). The inter-species and intra-species significance were obtained by one-way analysis of variance (one-way ANOVA). Directional selection: genes with significant differences between species and insignificant differences within species; stabilizing selection: genes with insignificant differences between species and within species; genetic drift: between species Genes with significant differences and intraspecies differences, and complex scenarios: Genes with insignificant differences between species and significant differences within species. Red: *Indica* rice strain; Blue: *Japonica* rice strain.

### The difference in expression level of directional selection genes and stabilizing selection genes

Species differentiation is affected by two types of genes: stabilizing selection and directional selection. Genes with small differences between species and within species when genes are subjected to stabilizing selection are the continuous selection pressure for interspecific traits in the population, and this natural selection will not lead to population differentiation (Charlesworth et al., 1982); genes subjected to directional selection are genes with large differences in expression between species and small differences in expression within species. They are the continuous strengthening of selective pressure for a trait among populations, and are the main genes that cause differences in traits between species (Mitchell-Olds et al., 2007; Rieseberg et al., 2002). Through previous research and analysis, we found that *indica* rice and *japonica* rice are mainly subject to two types of genes, namely stabilizing selection and directional selection, respectively. We analyze the differences in their expression levels between subspecies strains. We get TPM median of *indica* gene level is 7.3773, TPM median of gene subjected to stabilizing selection is 3.6448 . The expression level of directional selection genes in *indica* rice was significantly greater than that of stabilizing selection genes (Figure 3A Wilkerson rank sum test, p<0.001). In addition, the coefficient of variation (0.1364) of directional selection genes was significantly lower than that of stabilizing selection genes (0.2451) (Figure 3A Wilkerson rank sum test, p<0.001). We also found the same situation in *japonica* rice. The median TPM expression level of the directional selection gene in *japonica* rice was 9.4966, and the gene expression level of the stabilizing selection gene was 3.4431. The expression level of directional selection genes was significantly greater than that of stabilizing selection genes (Figure 3B Wilkerson rank sum test, p<0.001). Similarly, the coefficient of variation of *japonica* gene subjected to directional selection (0.1106) is significantly lower than the coefficient of variation of gene subjected to stabilizing selection (0.1904) (Figure 3B Weierkesen rank sum test, P <from 0.001). Therefore, from the data of gene expression level and variation level, we can conclude that the expression level of targeted selection genes is higher and the variation level is lower. Directional selection genes play a more important role in the process of subspecies differentiation than stabilizing selection genes.

**Figure 3.**
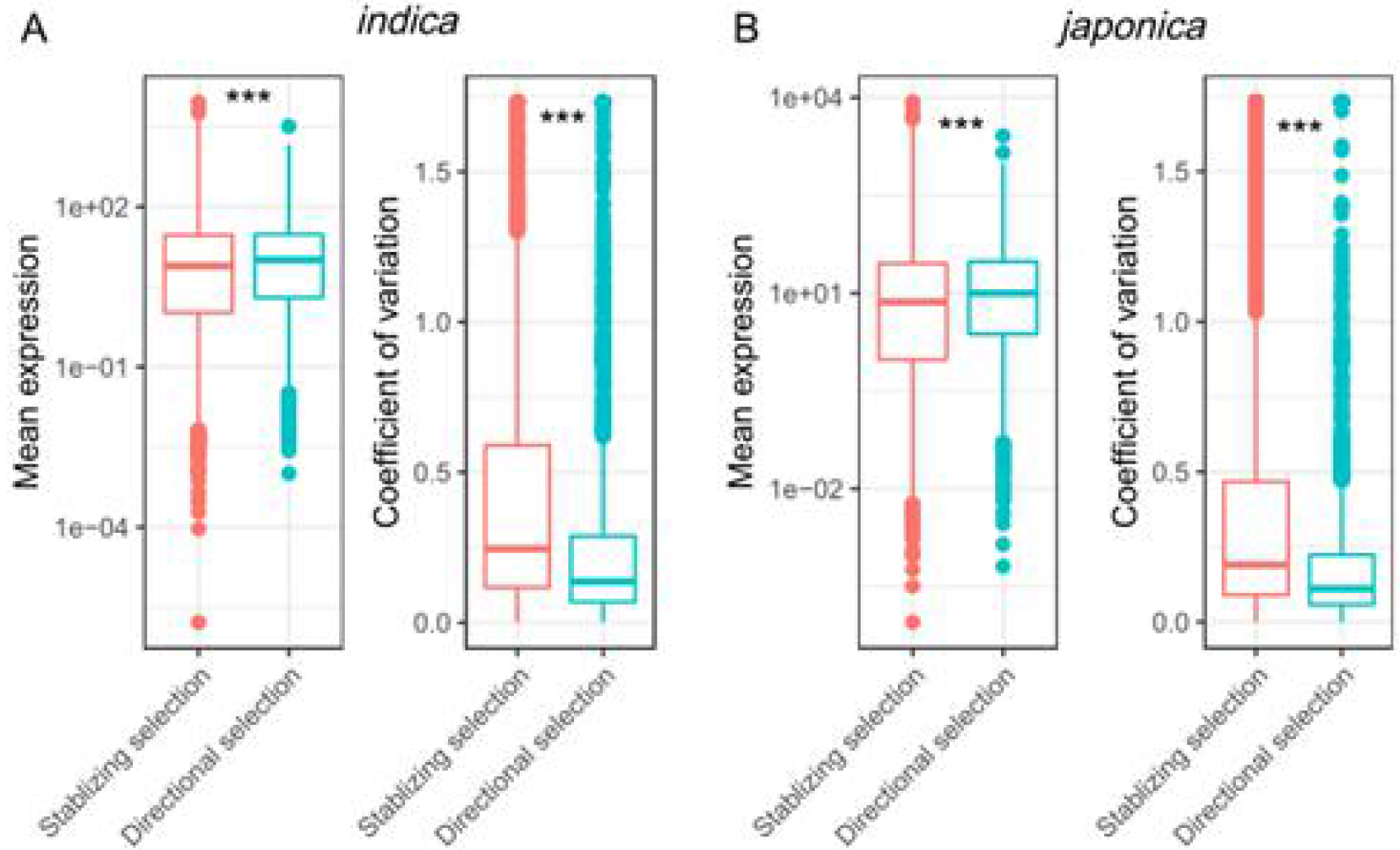
The expression and variation levels of directional selection and stabilizing selection in *indica* (A) and *japonica* (B). *** Wilcoxon rank sum test, p<0.001. Red: stabilizing selection gene; green: directional selection gene.

### *Indica* rice species-specific targeted selection genes will enrich biological pathways to cope with environmental pressures

In order to measure the phenotypic difference between *indica* and *japonica* in gene expression levels, we focused on species-specific directional selection of genes. Among the 5,910 *indica*-directional selection genes identified, 298 were *indica*-specific; among the 5,854 *japonica*-directional selection genes, 242 were *japonica*-specific. Subsequently, we will conduct gene function enrichment analysis for these subspecies-specific targeted selections (see Materials and Methods) to obtain functional pathways related to phenotypic differences. In addition to enriching a large number of common functional pathways, we found that *indica* rice-specific targeted genes also specifically enriched two types of biological pathways (Figure 4). The first category is related to coping with environmental stress, such as the response to stimulus, response to stress, response to abiotic stimulus (Figure 4A). This is related to the different geographical distribution, temperature and light adaptation of the two subspecies of *indica* and *japonica* (Foll & Gaggiotti, 2006; Mahgoub, 2019; Wu et al., 2019). The second category is the biosynthesis and metabolism, especially the Biopolymer Metabolism, Carbohydrate Metabolism, Protein biosynthesis, biosynthesis Macromolecule, etc (Figure 4A). This type of pathway may be related to the difference in starch and protein composition between *indica* and *japonica* rice. For example, indica rice has a higher amylose content than *japonica* rice (DuPont & Altenbach, 2003; Padilla-González et al., 2019). In addition, *indica* subspecies-specific targeted selection genes also have characteristics in the location of cellular components, for example, they are specifically located in non-membrane-bound organelles (intracellular non-membrane-bound organelle, external encapsulating structure). The molecule function also exist specificity, such as the activity of the specific structure of the molecules enriched (Structural Molecular Activity), transferase activity, etc.

**Figure 4.**
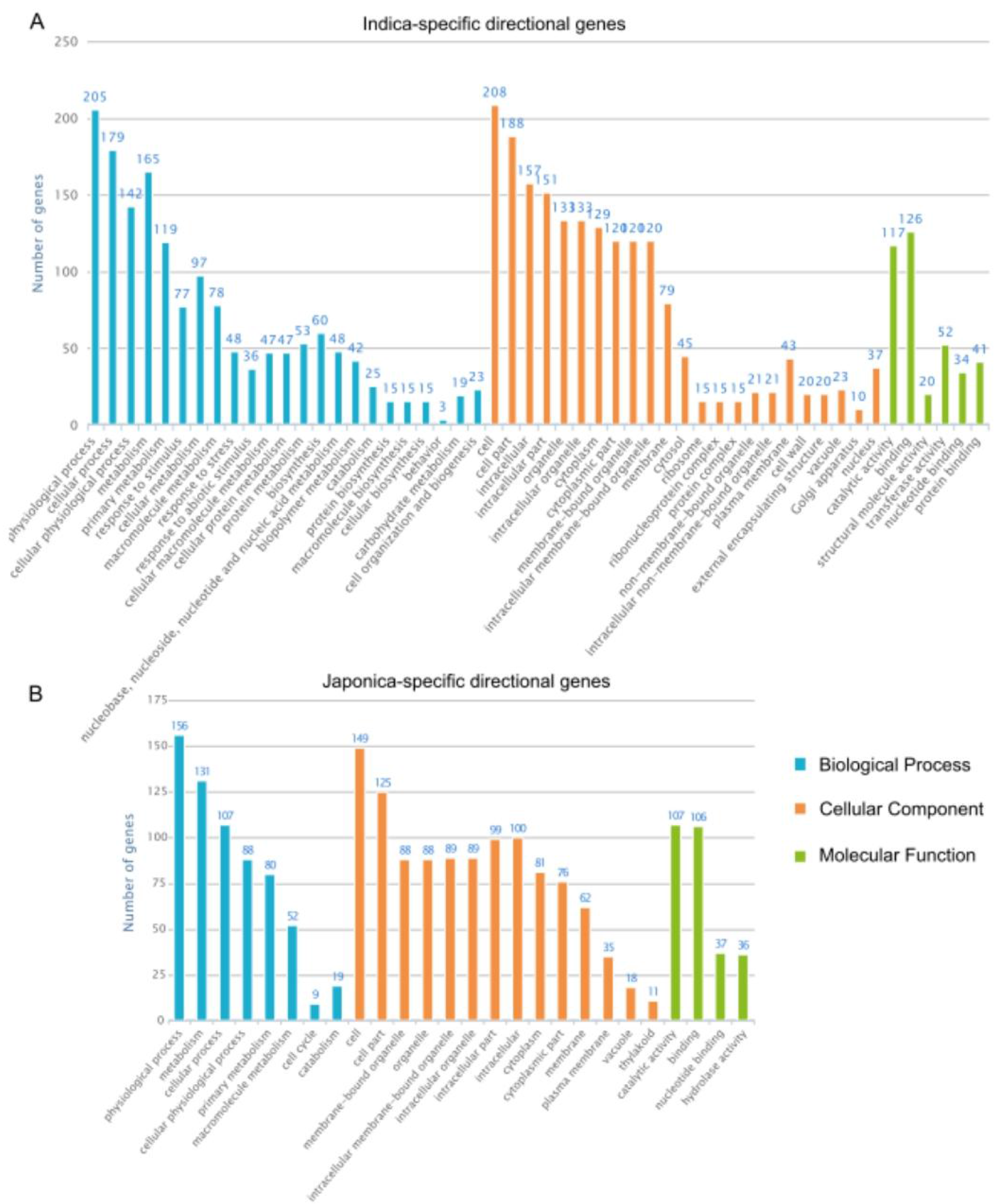
Gene Ontology (GO) functional enrichment analysis of species-specific directional selection genes in *indica* (A) and *japonica* (B). Blue: biological processes; orange: cellular components; green: molecular functions.

## Discussion

In the process of natural differentiation or artificial selection of species, the core gene regions affected by selection pressure are of concern to a wide range of researchers. If gene flow continues to affect species undergoing natural differentiation or artificial selection, under different selection pressures, different regions of the same genome will express different evolutionary results (He et al., 2011) , such as gene introgression. Differences in genomic DNA and changes in epigenetics are the two main factors that cause differences between species. Unlike in the past, which mainly focused on the change of DNA sequence to study species differentiation (He et al., 2011) , in recent years, more studies have focused on the transcriptome level and the expression level to explore the factors of species evolution. These studies have found that gene expression has a wide range of adaptability and is related to species differentiation (Guo et al., 2016; Wen et al., 2016; Yeh et al., 2014). In the past, some scholars studied the effects of changes in the species transcriptome on the evolution of gene expression, and gene chips. But our research has data advantages. We study the evolution of gene expression, which makes it easier to accurately find genes that differentiate between species than pure differential expression. In addition, we use more types of strains and effective methods to obtain more accurate transcriptomes (Ding et al., 2007; Goff et al., 2002; Huang et al., 2012; Wang et al., 2014) .

If the effects on rice differentiation are studied by observing changes in genome, this will have limitations and deficiencies, such as only studying wild rice and neglecting artificial selection (Guo et al., 2016) , or lack of measuring the internal expression variation of multiple strains between subspecies (Guo et al., 2016), or lack of sequencing data of tissue or developmental stages at that time (Wen et al., 2016). We use the linear model method to study the interspecific and intraspecific expression differences of the seedling transcriptome data of six lines of *japonica* rice and *indica* rice, and divide the genes into four selected gene types. We found that most of the subspecies with a higher degree of genome similarity are genes that are subjected to stabilizing selection, followed by genes that are subject to targeted selection. Later, in further comparing the expression levels and expression variation levels of genes subject to stabilizing selection and targeted selection, we found that targeted selection genes have higher expression levels and lower variation levels than stabilizing selection genes. This illustrates the importance of directional selection of genes in species differentiation. Finally, in the enrichment analysis of genes subject to targeted selection, we found that these genes are related to environmental pressure. This can indicate that the directional selection genes of *japonica* rice may have been affected by environmental pressure, which led to the differentiation of *indica* rice. The study of species evolution from the perspective of the transcriptome can provide different perspectives and depths.

Our research has two main shortcomings. First, from the data level, we only collected the transcriptome data of *indica* and *japonica* rice seedlings but missed the transcriptome data of other tissues or developmental stages. Future research can use more extensive and comprehensive organization and developmental stages to carry out more in-depth research. Second, there are many models for the evolution of cultivated rice. There are single-source models, that is they are domesticated together and then differentiated. There are also multiple independent domestication models, that is *indica* rice originates from *japonica* rice, and there are independent-origin models, that is completely independent domestication. In addition, wild rice, the ancestor of cultivated rice, is also controversial (He et al., 2011; Stein et al., 2018). However, we only studied the transcriptome of two cultivated rice. Future research can focus on the transcriptome of different wild rice. Although our results are insufficient, we have discovered the contribution of the transcriptome of cultivated rice seedlings to species differentiation. This lays the foundation for future research on the impact of changes in gene expression levels on species differentiation and gives certain prospective results, especially the impact of artificial selection on cultivated rice.

